# Classical drug digitoxin inhibits influenza cytokine storm, with implications for COVID-19 therapy

**DOI:** 10.1101/2020.04.09.034983

**Authors:** Bette S. Pollard, Jorge C. Blanco, John R. Pollard, Gregory A. Prince

**Affiliations:** Silver Pharmaceuticals, Rockville, MD, 20854; Sigmovir Biosystems, Rockville, MD, 20850; Department of Neurology, University of Pennsylvania, Philadelphia PA (USA) and Christiana Care Epilepsy Center, Newark, DE, 19173; Soft Cell Biological Research, St. George, UT, 84770

## Abstract

Influenza viruses, corona viruses and related pneumotropic viruses cause sickness and death partly by inducing a hyper-proinflammatory response by immune cells and cytokines in the host airway. Here we show that the cardiac glycoside digitoxin suppresses this response induced by influenza virus strain A/Wuhan/H3N2/359/95 in the cotton rat lung. The cytokines TNFα, GRO/KC, MIP2, MCP1, TGFβ, and IFNγ. are significantly and differentially reduced. Since the hyper-proinflammatory expression of cytokines is a host response, we suggest that digitoxin may have therapeutic potential for not only influenza and but also for coronavirus infections.

## Introduction

Influenza viruses, corona viruses and related pneumotropic viruses cause sickness and death partly by inducing a hyper-proinflammatory immune response in the host airway. This immune overreaction, called a cytokine storm, can lead to multiorgan failure and death (Tisoncik et al., 2012). For example, Influenza A (H5N1) has been shown to activate the TNFα-driven NFκB signaling pathway in a mouse host during viral infection generating a cytokine storm (Schmolke et al., 2009). As anticipated, inhibitors of NFκB acutely suppress cytokine storm and increase survival in a mouse model of SARS CoV infection (DeDiego et al., 2014). Recent data show that COVID-19 also activates NFκB (Guo et al., 2020). Cytokine storm marks the airways of SARS-CoV-2-infected patients that were admitted to the Intensive Care Unit (ICU) with more severe disease (Huang et al., 2020). Since there are multiple strains of influenza as well as coronavirus, there might be an advantage to develop therapies that suppress host-induced cytokine storm, in addition to developing strain-specific vaccines.

The clinical problem is that there are limited options for treating respiratory cytokine storm, most of which are predicated on inhibiting NFκB-activated cytokine expression (Yang et al., 2013a; Teijaro et al., 2014; Yang et al., 2017). The absence of NFκB inhibitory drugs from the human formulary is due to most candidate drugs being either neurotoxic or nephrotoxic when administered chronically (Zhang et al., 2017). One drug that lacks these toxicities is the cardiac glycoside digitoxin. We have previously shown digitoxin to be among the most potent inhibitors of the proinflammatory TNFα/NFκB pathway in the human airway and in other epithelial cells, both *in vitro* (Srivastava et al., 2004), and *in vivo* (Zeitlin et al., 2017; Pollard et al., 2019; Yang et al., 2019). Corroborating this is a screen of 2800 drugs and bioactive compounds which found digitoxin to be the 2nd most potent inhibitor of TNFα/NFκB activity (Miller et al., 2011). Digitoxin has been a drug to treat heart failure for decades, and is safe for children and adults with normal hearts (Hoffman and Bigger, 1990). In a clinical trial of digitoxin administered to young adults with the proinflammatory lung disease cystic fibrosis, digitoxin was safe. The study showed “the mRNAs encoding chemokine/cytokine or cell surface receptors in immune cells were decreased in nasal epithelial cells…leading to pathway-mediated reductions in IL-8, IL-6, lung epithelial inflammation, neutrophil recruitment and mucus hypersecretion.” (Zeitlin et al., 2017).

To further test the ability of digitoxin to inhibit cytokine storm related pneumotropic viruses, we used the cotton rat model of influenza infection to investigate the effects of digitoxin in influenza-induced cytokine storm. The cotton rat model has the important advantage of susceptibility to influenza infection without engineered adaptation. In addition, it has been shown that the response of the cotton rat to this virus strain evokes a pattern of pulmonary cytokine changes that parallel the human response (Ottolini et al., 2005).

## Results

### Digitoxin blocks cytokine storm

**Figure 1** shows the changes in cytokine protein in the lung due to digitoxin administration in the cotton rat after intranasally infected with 10^7^TCID50/100 gm of animal of influenza strain A/Wuhan/H3N2/359/95 virus. Animals were given three different doses of digitoxin, starting one day prior to virus administration and continuing with a daily dose until sacrifice on day 7. The maximum dose of digitoxin, 30 μg/kg, was calculated to be similar to the human dose routinely used to treat heart failure. As shown in **Figure 1**, protein data were collected for **IFNγ**(Interferon gamma); **GRO/KC**(rodent equivalent of human IL8); **MIP2**(Chemokine (C-X-C motif) ligand 2, CXCL2, Macrophage inflammatory protein 2-alpha); **TNFα**(Tumor Necrosis Factor alpha); **IL-1β**(Interleukin one beta); **MCP1**(Monocyte chemoattractant Protein 1, CCL2); and **TGFβ**(Transforming Growth Factor beta). As summarized in **Table 1**, digitoxin-dependent changes in protein were found to be significant for 6 of the 7 cytokines. The digitoxin-dependent reductions are specific and saturating for each cytokine, but do not reduce any of them to zero.

**Table 1.**
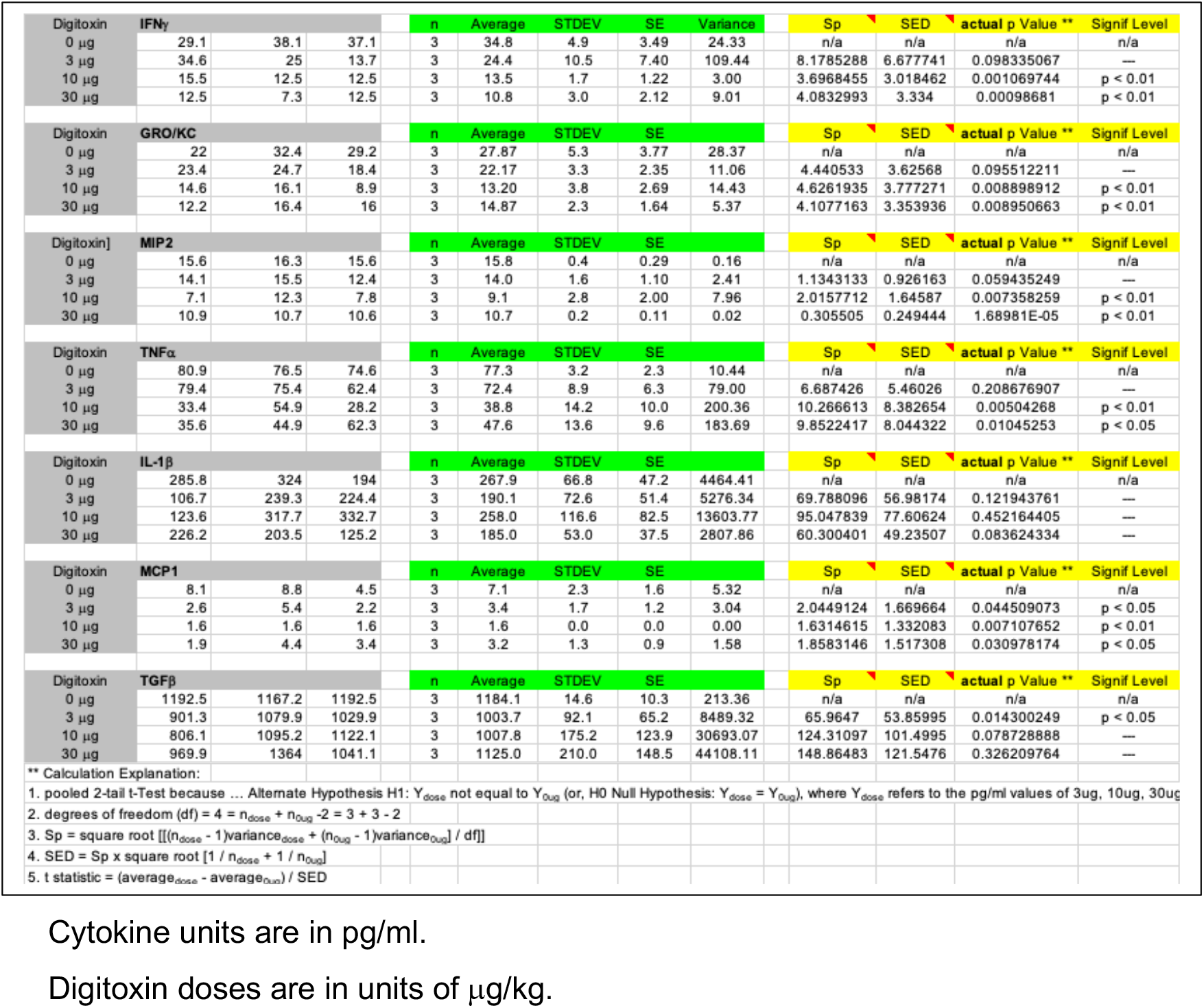
*Analysis of Digitoxin Suppression of Cytokine Expression in Cotton* Rat Lung Following Nasal Installation of Influenza Strain A/Wuhan/H3N2/359/95 Virus.

**Figure 1.**
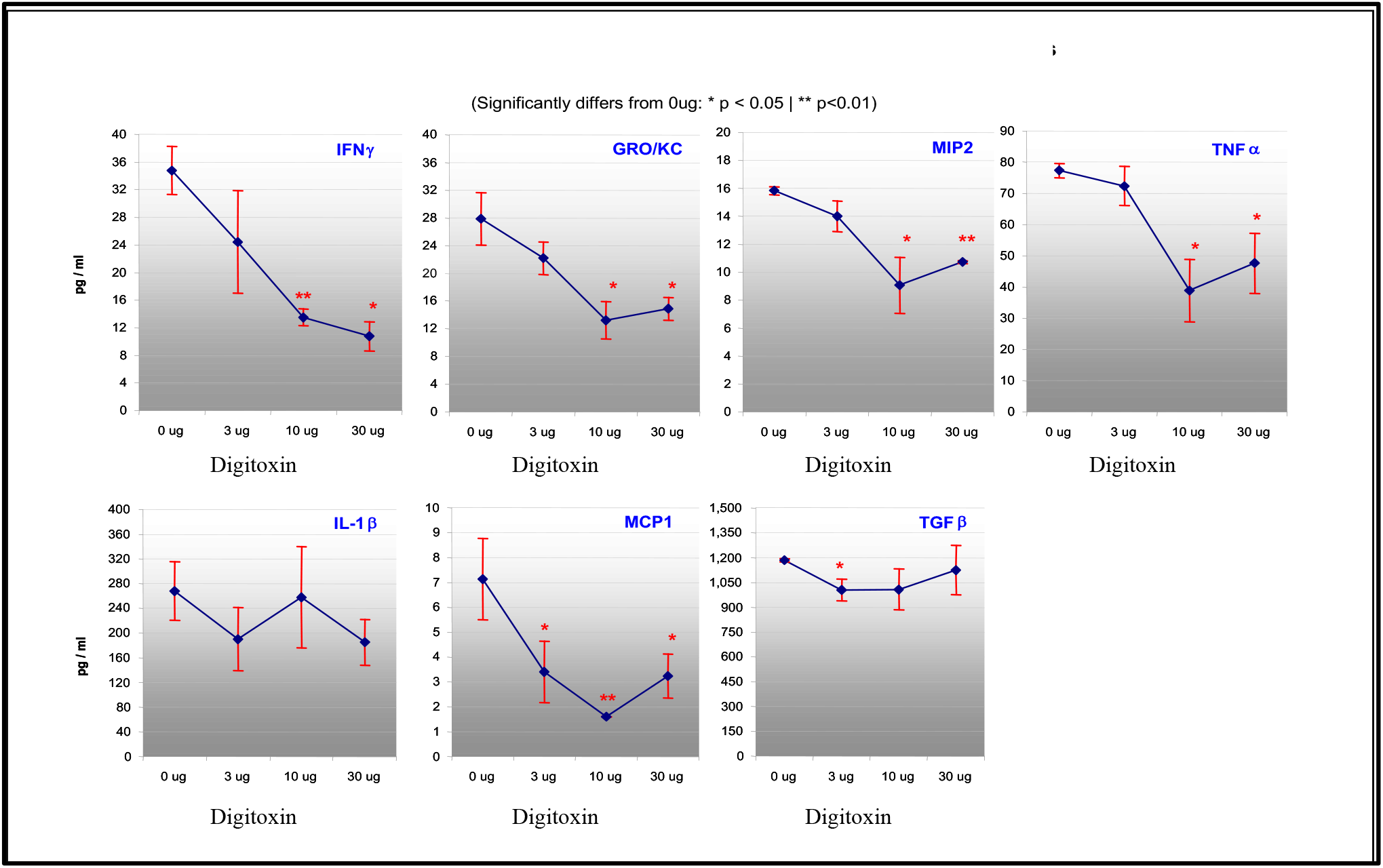
Cotton rats treated with digitoxin and influenza strain A/Wuhan/H3N2/359/95 virus. Animals were treated with different concentrations of digitoxin one day before intranasal virus administration and thereafter for 7 days. Samples assayed were lung tissue. Digitoxin dose is in units of μg/kg. Abbreviations are **INFγ**(interferon gamma, IFNγ); **GRO/KC**(Chemokine (C-X-C motif) ligand 1, CXCL1); **MIP2**(Chemokine (C-X-C motif) ligand 2, CXCL2, macrophage inflammatory protein 2-alpha); **TNFα**(TNFalpha, TNFA, tumor necrosis factor alpha); **IL-1β**(IL1B, interleukin 1 beta); **MCP1**(Monocyte chemoattractant Protein 1, CCL2); **TGFβ**(TGFB, transforming growth factor beta). Significance: * (*p* < 0.05); ** (*p* < 0.01); N = 3.

### Digitoxin differentially affects cytokine expression

**Table 1** shows that the greatest significant digitoxin-dependent reductions in cytokine proteins were found for **IFNγ** (68.9%), **GRO/KC**(46.6%), and **MCP1**(54.9%). Smaller but still significant reductions in cytokine proteins were found for **MIP2**(32.2%) and **TNFα** (38.4%). As also shown in **Table 1**, a significant reduction of only 15.3% was found for **TGFβ** cytokine protein at a concentration of 3 μg/kg, while only trending significance was noted at higher digitoxin concentrations. In the case of **IL-1β** only the highest digitoxin concentration trended towards significance. Thus digitoxin independently, dose-dependently and significantly lowers the individual concentrations in the lung of at least these six cytokines which have been induced by viral exposure.

## Discussion

These data show that administration of digitoxin to the cotton rat inhibits expression of many cytokines in the lung that are induced by influenza strain A/Wuhan/H3N2/359/95, including TNFα, the key activator of the TNFα/NFκB inflammation pathway. Digitoxin inhibits cytokine storm without compromising the entire immune system from physiologically responding to viral infection. The data also indicate that digitoxin acts on multiple cell types. For example, IFNγ is secreted only from activated T lymphocytes and NK cells of the immune system (Mah and Cooper, 2016). The remainder of the cytokines are secreted by epithelial cells in the airway, as well as by endothelial cells, immune cells and others (Mills et al., 1999; Liu et al., 2017). GRO/KC (CXCL1, the rodent equivalent of human IL8), a key target of NFκB signaling, is the most powerful known chemoattractant for drawing neutrophils into the lung. MIP2 and MCP1 induce entry and accumulation of monocytes and macrophages into the lung, and are targets of NFκB. TGFβ drives, and is driven by, NFκB-signaling for inflammation and fibrosis. IL-1β also drives NFκB, and is driven by NFκB. It appears that digitoxin-dependent reduction in TNFα/NFκB signaling is sufficient to suppress influenza A-driven cytokine storm.

In the cotton rat lung, IFNγmRNA expression in response to infection is biphasic (Ottolini et al., 2005). There is an early phase, from 6 hours after infection on day 1 to day 6, which may reflect the presence of activated NK cells. The late phase, from day 6 to day 28, may be the product of incoming antigen-specific T cells. Importantly, simply neutralizing INFγin a mouse model of infection with influenza A virus strain A/California/07/2009 (H1N1v; “Swine Flu”) is sufficient to not only alleviate acute lung injury but also to increase weight and survival rate (Liu et al., 2019). Reduced IFNγ is associated with reduced TNFα and NFκB activation. The data also show that digitoxin treatment causes the most profound reduction in INFγ expression. It is further known that IFNγ expression is driven by a combination of both NFκB and NFAT (Sica et al., 1997). As summarized in **Figure 2**, digitoxin not only reduces NFκB expression, but also reduces NFAT through digitoxin-dependent activation of Caspase 3 (Yang et al., 2013b).

**Figure 2.**
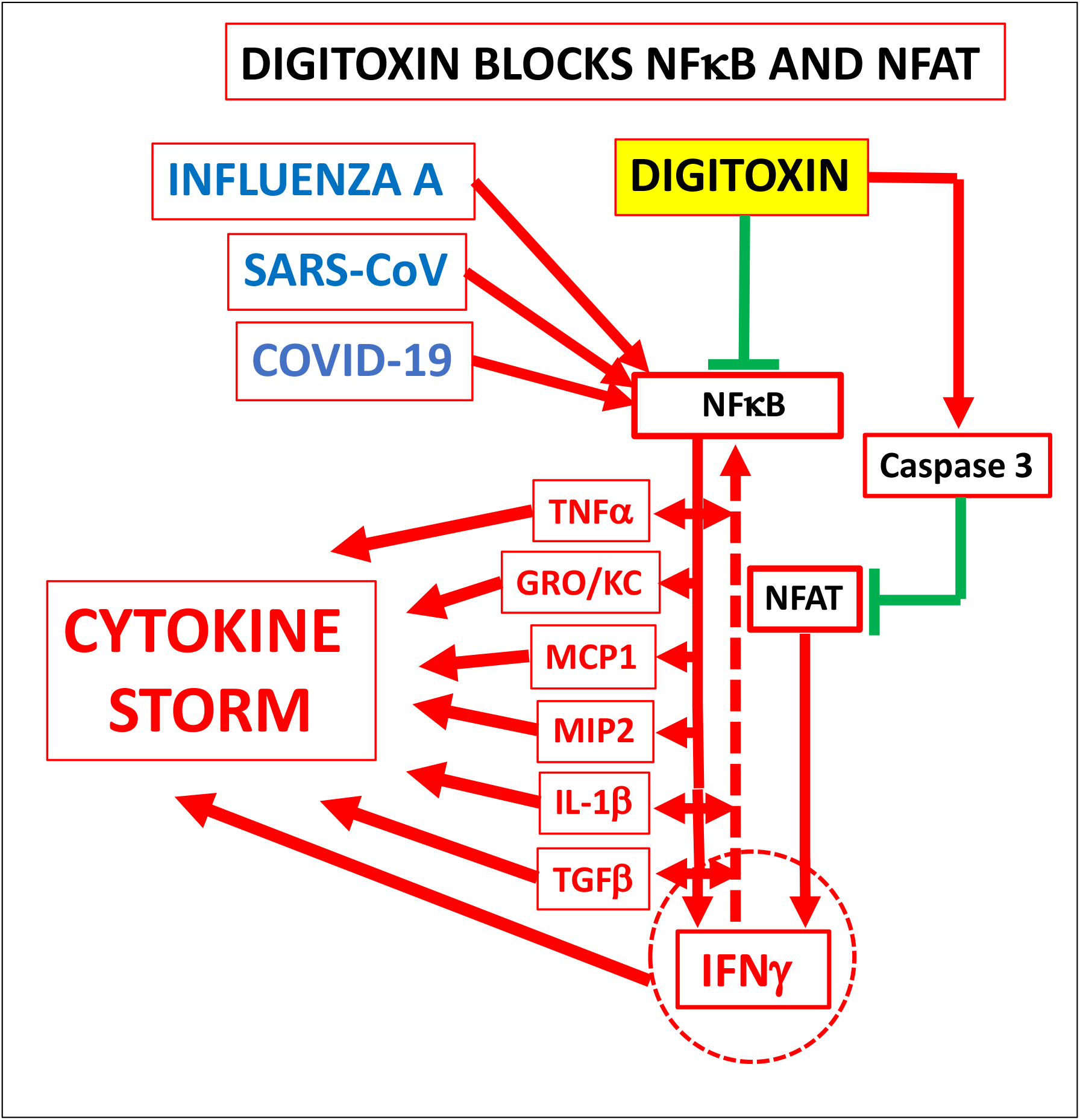
Digitoxin blocks cytokine storm, and interferon gamma. Influenza A virus, SARS-CoV and COVID-19 induce cytokine storm in the host. Present data show that digitoxin blocks cytokine storm when cotton rats are challenged with influenza strain A/Wuhan/H3N2/359/95. Expression of these cytokines and chemokines depend on NFκB, which is blocked by digitoxin. TNFα- IL-1βand TGFβalso activate NFκB. In addition, interferon gamma (INFγ) is blocked by digitoxin. INFγalso indirectly activates NFκB. Digitoxin activates caspase 3, which, proteolyzes NFAT. Experiments with SARS-CoV show that inhibition of NFκB suppresses cytokine storm and enhances survival in a mouse model (see text and DeDiego et al., 2014) COVID-19 activates NFκB (Guo et al., 2020) Cytokine storm induced by COVID-19 has been observed in severely affected patients (Huang et al., 2020). Color code: red (activation); green (inhibition); See cytokine abbreviations in Figure 1. Dotted line represents intervening steps.

Finally, since antiviral properties have been reported for digitoxin and other cardiac glycosides, it is a limitation of the study that we cannot exclude other antiviral effects by digitoxin from contributing to the reduction in influenza-dependent cytokine concentrations (Burkard et al., 2015; Amarelle and Lecuona, 2018; Wei et al., 2020).

In conclusion, these data show that digitoxin blocks the host cytokine storm induced by influenza strain A/Wuhan/H3N2/359/95 in the cotton rat lung. Since digitoxin already had been shown to be safe in CF patients with pulmonary disease and a normal heart, and caused a similar reduction in NF-KB driven cytokine expression, this drug may be a good candidate for further investigation as a therapy for influenza and potentially for COVID-19.

## Methods

### Animal protocol

Cotton rat experiments were performed as previously described (Ottolini et al., 2005). All experiments were performed using protocols that followed federal guidelines and were approved by the Institutional Animal Care and Use Committee. Animals were sacrificed by carbon dioxide inhalation.

### Drugs and protocol for drug preparation

Digitoxin (μg/kg) was obtained from Sigma-Aldrich (> 95% pure). The drug was prepared as a stock solution in 95% ethanol, and further diluted in PBS before administration. Digitoxin was administered to cotton rats intraperitoneally one day before intranasal infection with 10^7^TCID50/100 gm cotton rat with influenza strain A/Wuhan/H3N2/359/95. Daily digitoxin treatment continued until harvest on day 7 of the experiment.

### Tissue preparation and histological analysis

On day 7 of the experiment, the animals were sacrificed. The left lung was first tied off and reserved for cytokine analysis. Lung samples were then immediately frozen on dry ice, and then kept at −80°C until further processed. The remaining lung tissue was processed for histological analysis. Histology revealed no apparent change due to administration of the drug.. The frozen tissues were transferred to Silver Pharmaceuticals.

### Biochemical analysis

Frozen lung samples were weighed, thawed and then minced with scissors in 10% (w/v) ice cold PBS, homogenized in 10 strokes in a Ten Broeck homogenizer, and centrifuged at 20,000 X g for 30 minutes. The supernatant solutions were kept at −80°C until assay. The supernatant solutions were tested by Silver Pharmaceuticals for cytokines and chemokines by ELISA at Bioassay Works, LLC in Ijamsville, MD. The samples were then sent for corroboration to Pierce-Thermo for ELISA assay on the Searchlight® ELISA platform. Rat antibody reagents were used in both instances.

## Acknowledgements

The authors thank Val Hemming, M.D. and Harvey B. Pollard, M.D., Ph.D. for advice, and Mr. Max Tran, MBA, for providing statistical analysis.

## Notes

### Competing Interest Statement

The authors have declared no competing interest.

